# Waning immune responses against SARS-CoV-2 among vaccinees in Hong Kong

**DOI:** 10.1101/2021.12.22.473934

**Authors:** Qiaoli Peng, Runhong Zhou, Yuewen Wang, Meiqing Zhao, Na Liu, Shuang Li, Haode Huang, Dawei Yang, Ka-Kit Au, Hui Wang, Kwan Man, Kwok-Yung Yuen, Zhiwei Chen

## Abstract

**Background:** Nearly 4 billion doses of the BioNTech-mRNA and Sinovac-inactivated vaccines have been administrated globally, yet different vaccine-induced immunity against SARS-CoV-2 variants of concern (VOCs) remain incompletely investigated.

**Methods:** We compare the immunogenicity and durability of these two vaccines among fully vaccinated Hong Kong people.

**Findings:** Standard BioNTech and Sinovac vaccinations were tolerated and induced neutralizing antibody (NAb) (100% and 85.7%) and spike-specific CD4 T cell responses (96.7% and 82.1%), respectively. The geometric mean NAb IC_50_ and median frequencies of reactive CD4 subsets were consistently lower among Sinovac-vaccinees than BioNTech-vaccinees. Against VOCs, NAb response rate and geometric mean IC_50_ against B1.351 and B.1.617.2 were significantly lower for Sinovac (14.3%, 15 and 50%, 23.2) than BioNTech (79.4%, 107 and 94.1%, 131). Three months after vaccinations, NAbs to VOCs dropped near to detection limit, along with waning memory T cell responses, mainly among Sinovac-vaccinees.

**Interpretation:** Our results indicate that Sinovac-vaccinees may face higher risk to pandemic VOCs breakthrough infection.

**Funding:** This study was supported by the Hong Kong Research Grants Council Collaborative Research Fund (C7156-20GF to Z.C and C1134-20GF); the National Program on Key Research Project of China (Grant 2020YFC0860600, 2020YFA0707500 and 2020YFA0707504); Shenzhen Science and Technology Program (JSGG20200225151410198 and JCYJ20210324131610027); HKU Development Fund and LKS Faculty of Medicine Matching Fund to AIDS Institute; Hong Kong Innovation and Technology Fund, Innovation and Technology Commission and generous donation from the Friends of Hope Education Fund. Z.C.’s team was also partly supported by the Theme-Based Research Scheme (T11-706/18-N).

## Introduction

Severe acute respiratory syndrome coronavirus 2 (SARS-CoV-2) is the causative agent of coronavirus disease 2019 (COVID-19). SARS-CoV-2 has been spreading worldwide since December 2019, leading to the ongoing COVID-19 pandemic with 234 million infections and 4.8 million deaths by 30 September 2021 (https://covid19.who.int/). Due to pressure of the COVID-19 pandemic, the greatest global efforts have been placed for vaccine development. To date, six vaccines have been approved by regulatory agencies for emergency use including (1) two mRNA-based vaccines, namely BNT162b2 (by Pfizer Inc. and BioNTech SE) and mRNA-1273 (by Moderna), expressing full spike (S) glycoprotein with efficacy rates of 94.1-95% (1, 2), (2) the chimpanzee adenovirus-vectored vaccine, named ChAdOx1 nCoV-19 (by the Oxford University and AstraZeneca Inc.), encoding the full S glycoprotein with an efficacy rate of 70.4% (3), (3) the human adenovirus-vectored vaccine, namely Ad26.COV2.S (by Johnson & Johnson Inc.), encoding the full S glycoprotein with an efficacy rate of 73.1% (4), (4) two inactivated vaccines CoronaVac and BIBP (by Sinovac Biotech and SinoPharm) with efficacy rates of 83.7% (5) and 78.1% (6), respectively. In recent reports, however, SARS-CoV-2 variants of concerns (VOCs) have posted challenges for vaccine-induced protection (7–9).

Over four million genome sequences of SARS-CoV-2 have been submitted to the hCoV-19 database of the Global initiative on sharing all influenza data (GISAID) since the outbreak of COVID-19. Several VOCs have had significant impacts on the trend of the pandemic. The top five noticeable VOCs include B.1.1.7 variant (Alpha, United Kingdom), B.1.351 (Beta, South Africa), P1 (Gamma, Japan/Brazil), B.1.617.2 (Delta, India), and B.1.427/B.1.429 (Epsilon, United States) (10). The VOC B.1.351 strain was significantly resistant (10.3-12.4-fold) to neutralization by sera derived from vaccinated individuals who received Moderna or BioNTech compared to the VOC D614G strain (7). Although vaccinations reduced sickness, hospitalization and death rates, vaccine-induced attenuation of peak viral burden has decreased for the VOC B.1.617.2 strain compared to the VOC B.1.1.7 variant in the UK (11). These results are in line with the increasing number of breakthrough infections among fully vaccinated population (12). It is, therefore, critical to study vaccinees to determine their potential risk to the spreading VOCs.

Four waves of severe acute respiratory syndrome coronavirus 2 (SARS-CoV-2) epidemic have hit Hong Kong, resulting in 10950 infections and 198 deaths. To control the epidemic effectively, the BioNTech [COMIRNATY^™^] and Sinovac [CoronaVac] vaccines have been made available for Hong Kong residents since 26 February 2021 (www.covidvaccine.gov.hk/en/). In mainland China, more than 2.2 billion doses of inactivated vaccines, including mainly Sinovac, have been inoculated by 30 September 2021. Since both vaccines have been recommended by World Health Organization for emergency use(1) (5), 66.8% of the 7.5 million Hong Kong people have been fully vaccinated by 30 September 2021. However, the immunogenicity and durability of these two vaccines in terms of antibody and T cell responses against VOCs remain largely unknown. With the accumulating emergence and spreading of pandemic VOCs, monitoring vaccine-induced neutralizing antibody (NAb) and T cell memory responses, especially NAbs activity against VOCs, may play a critical role in determining the policy of boost vaccination. For this reason, we aimed to determine humoral and cellular immune responses in parallel among Hong Kong vaccinees over time with focus on cross-reactive NAbs against VOCs.

## Methods

### Study subjects

Participants who completed two doses of SARS-CoV-2 vaccination (either BioNTech or Sinovac) before June 2021 were recruited for this study. The exclusion criteria include individuals with (1) documented SARS-CoV-2 infection, (2) high-risk infection history within 14 days before vaccination, (3) COVID-19 symptoms such as sore throat, fever, cough and shortness of breath. This study was approved by the Institutional Review Board of the University of Hong Kong/Hospital Authority Hong Kong West Cluster (Ref No. UW 21-452). Written informed consent was obtained from all study subjects. Peripheral blood mononuclear cells (PBMCs) and inactivated plasma were freshly isolated for testing. Participants were required to record any adverse reactions related to each dose of vaccination, including local adverse reactions surrounding the injection site and systemic adverse reactions. The demographic characteristics of these two groups were similar in terms of gender, age, nationality, body mass index, etc. (Table 1). Besides participants with Chinese nationality, there were one Sinovac recipient from Malaysia and three non-Asian BioNTech recipients from France, America and Danish, respectively. Two participants in the BioNTech group have mild hypertension and diabetes. The median intervals between two doses and the time of blood collection after second dose were also comparable. To assess the immunogenicity of both vaccines, 16 gender- and age-matched non-vaccinated subjects were used as controls. These healthy donors did not have prior history of SARS-CoV-2 infection.

**Table1.** Demographic characteristics of study subjects

|  | BioNTech (n=34) | Sinovac (n=28) | *P* value |
| --- | --- | --- | --- |
| Gender, male (%) | 16 (47.1%) | 14 (50.0%) | 0.818 |
| Age, median years (IQR) | 30.5 (26.8-35.3) | 29.0 (26.0-31.0) | 0.085 |
| Nationality |  |  | 0.62 |
| Chinese | 31 (91.2%) | 27 (96.4%) |  |
| Non-Chinese | 3 (8.8%) | 1 (3.6%) |  |
| Place of vaccination |  |  | 0.345 |
| Hong Kong | 33 (97.1%) | 24 (85.7%) |  |
| Mainland | 0 (0.0%) | 4 (14.3%) |  |
| Others | 1 (2.9%) | 0 (0.0%) |  |
| BMI (Kg/m<sup>2</sup>) (IQR) | 21.3 (19.2-24.0) | 21.0 (18.8-26.6) | 0.253 |
| Underlying diseases |  |  | 0.497 |
| Yes | 2 (5.9%) | 0 (0.0%) |  |
| No | 32 (94.1%) | 28 (100.0%) |  |
| The median interval days between two vaccinations (IQR) | 30 (22-32) | 28 (28-30) | 0.493 |
| The median interval days between the second dose and the first blood collection (IQR) | 30 (22-32) | 28 (20-39) | 0.858 |

### Enzyme-linked immunosorbent assays (ELISA)

ELISA was used for determining the IgG binding to RBD and full spike as we previously described (13). The area under the curve (AUC), representing the total peak area based on ELISA OD values as previously described (14), of each sample was plotted using the GraphPad Prism v8, and the baseline with the defined endpoint was set as the average of negative control wells+10 standard deviation). The limit of quantification (LOQ) was established based on the geometric mean of non-vaccinated donors without prior SARS-CoV-2 infection history.

### Pseudotyped viral neutralization assay

The S-expression plasmids encoding wildtype, D614G, B.1.1.7, B.1.351, P1, B.1.617.2 and B.1.429 variants were used to generate pseudoviruses. P1 and B.1.429 were purchased from InvivoGen while others were made by us or collaborators. Briefly, SARS-CoV-2 pseudoviruses were generated by co-transfection of 293T cells with a pair of plasmids, the S-expression plasmid for wildtype or VOCs and the pNL4-3Luc_Env_Vpr plasmid in a human immunodeficiency virus type 1 backbone (15, 16). At 48 hours post-transfection, virus-containing supernatant was collected, quantified and frozen at −150°C. The pseudotyped neutralization assay of vaccinated samples was performed as previously described (15, 16). Serially diluted and heat-inactivated plasma samples were incubated with 200 TCID50 of pseudovirus at 37°C for 1 hour. The plasma-virus mixtures were then added into pre-seeded HEK 293T-hACE2 cells. After 48 hours, infected cells were lysed, and luciferase activity was measured using the Luciferase Assay System kit (Promega) in a Victor3-1420 Multilabel Counter (PerkinElmer). The 50% inhibitory concentrations (IC_50_) of each specimen were calculated using non-linear regression in GraphPad Prism v8 to reflect anti-SARS-CoV-2 antibody potency. Samples that failed to reach 50% inhibition at the lowest serum dilution of 1:20 were considered to be non-neutralizing, and the IC_50_ values were set to 10.

### Peptide pools

We purchased peptide pool of 15 amino acid (aa) overlapping by 11 aa spanning the full length of SARS-CoV2-spike (a total of 316 peptides), receptor binding domain (RBD)(S306-S543, a total of 57 peptides) and nucleocapsid protein (NP) (a total of 102 peptides) from GenScipt. As a control, we utilized a peptide pool spanning the entire region pp65 protein (15-mers overlapping by 11 aa) of human cytomegalovirus (CMV), which was obtained from the NIH HIV reagent program (CAT# ARP-11549).

### Intracellular cytokine staining (ICS)

To measure antigen-specific T cell response, PBMCs were stimulated with 2 μg/mL of indicated COVID-19 antigen peptide pools (RBD or Spike or NP) in the presence of 0.5 μg/mL anti-CD28 and anti-CD49d mAbs (Biolegend). Cells were incubated at 37 □ overnight, and BFA (Sigma) was added at 2 h post incubation, as previously described (13). CMV (pp65) peptide pool was included as an internal positive control. Stimulation alone with anti-CD28 and anti-CD49d was used as negative control. After overnight incubation, cells were washed with staining buffer (PBS containing 2% FBS) and stained with mAbs against surface markers (Zombie Aqua, Pacific blue anti-CD3, Percp-Cy5.5 anti-CD4, APC-Fire750 anti-CD8, BV711 anti-CD45RA and APC anti-CCR7) (Biolegend). For intracellular staining, cells were fixed and permeabilized with BD Cytofix/Cytoperm (BD Biosciences) prior to staining with the mAbs against cytokines (PE anti-IFN-γ, AF488 anti-TNF-α and PE-Cy7 anti-IL-2) (Biolegend) with Perm/Wash buffer (BD Biosciences). After gating on CD4^+^ T and CD8^+^ T cells, intracellular IFN-γ/TNF-α/IL-2 were calculated (Fig. S1). All percentages of antigen-specific CD4^+^ and CD8^+^ T cells were reported as background subtracted data from the same sample stimulated with negative control (anti-CD28/CD49d only). The LOQ for antigen-specific CD4^+^ (0.01%) and CD8^+^ T cell responses (0.02%) was calculated using a twofold median value of all negative controls. Responses >LOQ and a stimulation index >2 for CD4^+^ and CD8^+^ T cells were considered positive responder. Values higher than the threshold of positive responders after spike peptide pool stimulation were considered for the analysis of multifunctional antigen-specific T cell responses. Phenotype profiles were further analyzed by gating on IFN-γ^+^CD4 or IFN-γ^+^CD8 T cells for expression of CCR7 and/or CD45RA in response to spike, respectively.

### Statistical analysis

Flow cytometric data were analysed using FlowJo 10.6.0. Statistical analysis was performed using the GraphPad Prism v8 Software. Mann-Whitney U test was used to compare between-group continuous values. Wilcoxon signed-rank test was used for paired comparisons. For between-group categorical values comparison, two-sided chi-square tests or fisher’s exact test were used. The non-parametric Spearman test was used for correlation analysis. The statistical method of aggregation used for the analysis of binding and neutralizing antibody titers (NAbTs) is geometric mean titer (GMT) with the corresponding 95% confidence intervals (95%CI), and median with interquartile (IQR) for antigen-specific T cell frequencies. The statistic details are depicted in the respective legends. *P* < 0.05 was considered statistically significant.

## Results

### Safety profiles of BioNTech and Sinovac in Hong Kong vaccinees

A total of 62 vaccinated subjects were enrolled in this study including 34 BioNTech-vaccinees and 28 Sinovac-vaccinees. Blood samples were collected at acute phase with a median of 30 [IQR, 22 to 32] days for BioNTech and 28 [IQR, 20 to 39] days for Sinovac post the second dose. Samples from 27 BioNTech-vaccinees and 16 Sinovac-vaccinees were successfully followed up at memory phase with a median of 113 [IQR, 101 to 115] days and 105 [IQR, 96 to 109] days post the second dose, respectively. After the first vaccination, the overall incidence rate of any adverse reactions was higher for BioNTech than Sinovac (79.4% [27/34] versus 53.6% [15/28], *P*=0.03). The most common adverse reaction was the injection-site pain (73.5% [25/34] versus 35.7% [10/28], *P*=0.003), followed by fatigue (41.2% [14/34] versus 25.0% [7/28], *P*=0.18) and myalgia (17.6% [6/34] versus 14.3% [4/28], *P*=0.991), respectively (Fig. 1A). After the second vaccination, the overall incidence of any adverse reactions was consistently higher for BioNTech than Sinovac (88.2% [30/34] versus 57.1% [16/28], *P*=0.005), including the major injection-site pain (64.7% [22/34] versus 53.6% [15/28], *P*=0.374) and other systematic symptoms (e.g., headache, fever and fatigue) (Fig. 1B). Moreover, significantly more BioNTech vaccinees had more than two adverse reactions (86.7% [26/30] versus 31.3% [5/16], *P*<0.001). Most of the adverse reactions, however, were considered tolerable with an average recovery time of 48-72 hours. No vaccine-associated severe adverse reactions were reported among our study subjects.

**Figure 1.**
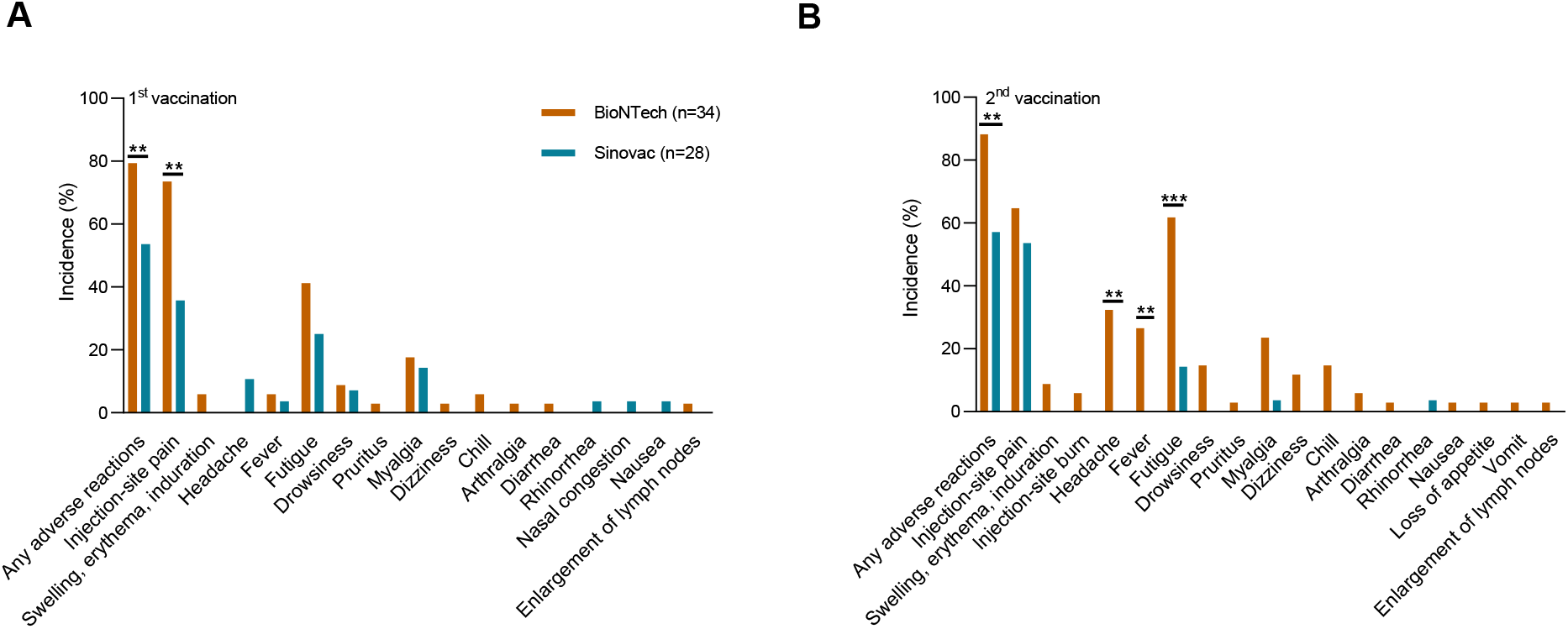
Adverse reactions after the first and the second vaccination of BioNTech and Sinovac. The incidence rates of adverse reactions were compared between BioNTech (orange) and Sinovac (green) after the first (A) and the second (B) vaccination. Data were analyzed for statistical significance using the two-sided chi-square test or the fisher’s exact test. \*\**P*<0.01, \*\*\**P*<0.001.

### Humoral and cellular immune responses induced by BioNTech and Sinovac

The amounts of anti-spike and receptor-binding domain (RBD) IgG as well as pseudovirus NAbTs were firstly determined at the acute phase after vaccination. Anti-spike and RBD IgG were induced among 100% vaccinees in the BioNTech group. In contrast, only 85.7% (24/28) of Sinovac vaccinees showed a detectable amount of anti-spike and RBD IgG. Compared to Sinovac, BioNTech induced a significantly higher GMT of anti-spike IgG (1400 [95CI% 1035 to 1894] versus 217.8 [95CI% 152.7 to 310.5], *P*<<0.0001) and anti-RBD IgG [683.3 (95CI% 498.4 to 936.9) versus 17.8 (95CI% 4.1 to 76.7), *P*<0.0001] responses (Fig. 2A). Encouragingly, the majority of BioNTech and Sinovac vaccinees (100% [34/34] versus 85.7% [24/28]) developed NAb responses against the wildtype Wuhan pseudovirus. Moreover, immunized sera from BioNTech vaccinees showed 19 times higher NAbTs against wildtype than that from Sinovac vaccinees based on the geometric mean IC_50_ (1401[95CI% 1076 to1823] versus 73.7 [95CI% 43.4 to 125], *P*<0.0001) (Fig. 2B). Since the neutralization potency index calculated by the NT_50_/IgG ratio was suggested as a predictor of survival (17), we also evaluated this factor by calculating the ratio of IC_50_ to AUC of anti-spike and RBD IgG (Fig. 2C). The neutralization potency index of BioNTech (geometric mean of 1.0 [95% CI 0.73 to 1.37] for IC_50_/spike IgG and 0.88 [95% CI 0.66 to 1.17] for IC_50_/RBD IgG) was significantly higher than that of Sinovac (0.36 [95% CI 0.24 to 0.55] for IC_50_/spike IgG and 0.18 [95% CI 0.09 to 0.35] for IC_50_/RBD IgG) (both *P* values <0.0001). These results demonstrated that while anti-spike IgG, anti-RBD IgG and NAbs were induced by both vaccines, the NAbTs of Sinovac vaccinees was 19-fold lower than that of BioNTech vaccinees, together with the lower neutralization potency index.

**Figure 2.**
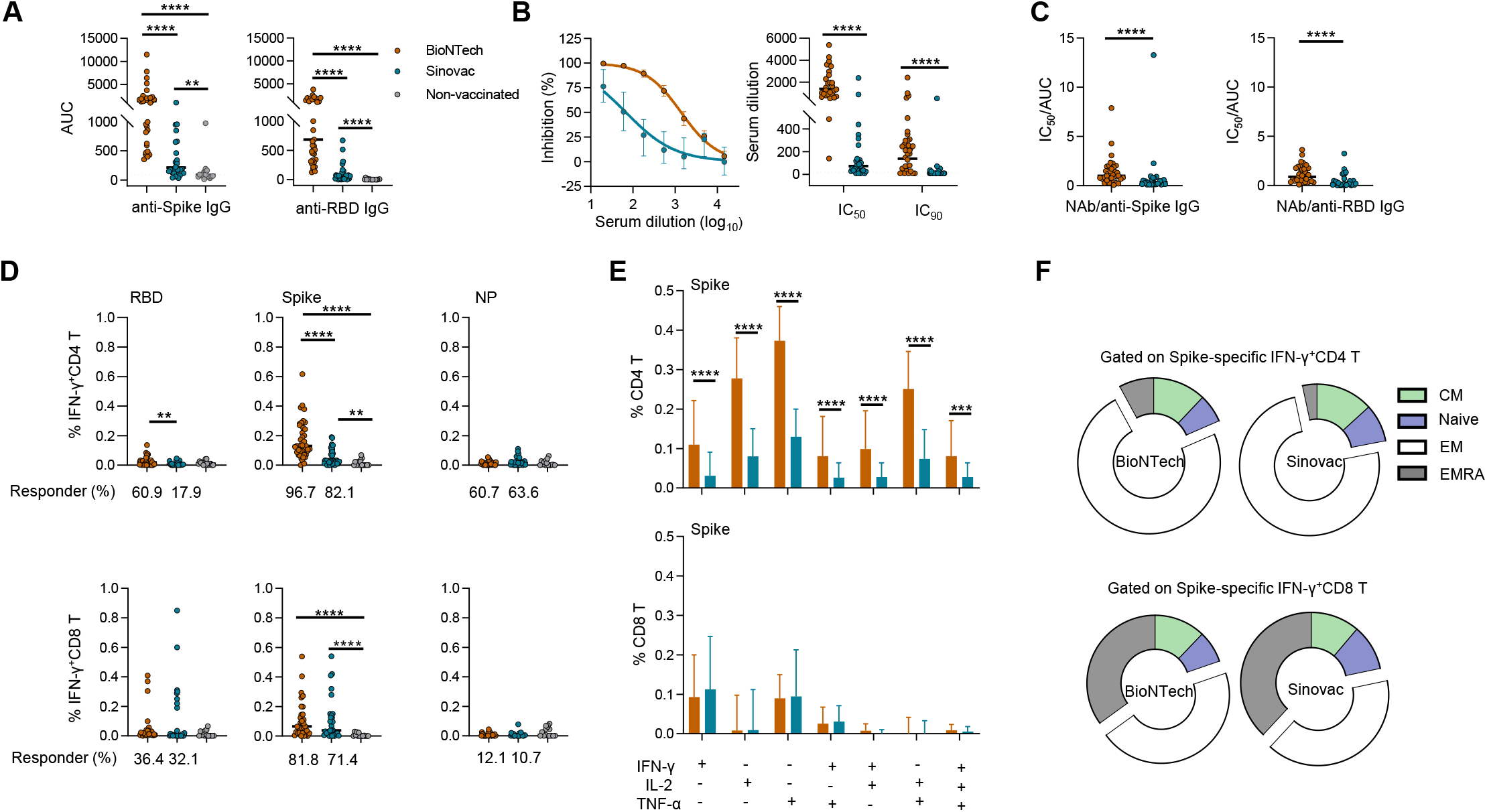
Immune responses measured at the acute phase after the second dose among BioNTech and Sinovac vaccinees. (**A**) The area under curve (AUC) of anti-Spike and anti-RBD IgG in BioNTech (orange) (n=34), Sinovac (green) (n=28) and non-vaccinated volunteers (grey)(n=16). The AUC represents the total peak area calculated from ELISA OD values by the GraphPad Prism v8. (**B**) Precent inhibition and IC_50_/IC_90_ values against wild type SARS-CoV-2 pseudoviruses in BioNTech and Sinovac vaccinees. (**C**) The neutralization antibody potency index defined by the ratio of IC_50_/AUC of anti-Spike IgG and anti-RBD IgG in BioNTech and Sinovac group. Data showed geometric mean values in each group in A-C. (**D**) Quantified results depict the percentage of RBD, spike and NP-specific IFN-γ^+^ CD4^+^ T (top) and IFN-γ^+^ CD8^+^ T (bottom) cells in BioNTech (n=33), Sinovac (n=28) and non-vaccinated volunteers (n=15), respectively. Fresh PBMC were subjected to T cell response measurement by ICS after RBD-, spike- and NP-specific *ex vivo* peptide pool stimulation, respectively. (**E**) The proportions of spike-specific polyfunctional CD4^+^ T (top) and CD8^+^ T (bottom) cells were compared in BioNTech and Sinovac-vaccinated responders. After gating on CD4 or CD8^+^ T cells, single cytokine (IFN-γ^+^ or TNF-α^+^ or IL-2^+^), double cytokines (IFN-γ^+^ TNF-α^+^ or IFN-γ^+^ IL-2^+^ or TNF-α^+^ IL-2^+^), and triple cytokines (IFN-γ^+^ TNF-α^+^ IL-2^+^) producing cells were analyzed in response to spike-specific *ex vivo* peptide pool stimulation, respectively. Background-subtracted data was analyzed in all cases in D and E. The bars in D and E indicated median with interquartile (IQR). (**F**) Phenotypic analysis depicted antigen-specific T cell subsets of BioNTech and Sinovac vaccinees. After gating on IFN-γ^+^ CD4^+^ or IFN-γ^+^ CD8^+^ T cells, T cell subsets expressing CCR7 and/or CD45RA were analyzed in response to spike-specific *ex vivo* peptide pool stimulation. Data were analyzed for statistical significance using Mann-Whitney U test. Dotted black lines indicate the limit of quantification (LOQ). \*\**P*<0.01, \*\*\**P*<0.001, \*\*\*\**P*<0.0001.

Besides humoral immune response, we also measured antigen-specific T cell response because it may play an important role in protection against SARS-CoV-2 infection (13, 18). Vaccine-specific T cell responses were determined by ICS after stimulation by the peptide pools covering RBD, spike and NP antigen (Fig. S1). Spike-specific IFN-γ^+^CD4^+^T cells were induced in 96.7% (32/33) of BioNTech and 82.1% (23/28) of Sinovac subjects. The frequencies of spike-specific IFN-γ^+^CD4^+^T cells in both vaccinated groups were significantly higher than those of the non-vaccinated controls. The median frequency was 0.13 [IQR 0.08 to 0.26] for BioNTech and 0.03 [IQR 0.02 to 0.08] for Sinovac as compared with controls’ 0.00 [IQR 0.00 to 0.02] (*P*<0.0001 and *P*=0.0017), respectively (Fig. 2D, top). Meantime, both BioNTech (81.8% [27/33]) and Sinovac (71.4% [20/28]) vaccinees displayed higher frequencies of spike-specific IFN-γ^+^CD8^+^T cells than the non-vaccinated group (0.07 [IQR 0.02 to 0.17] for BioNTech and 0.04 [IQR 0.02 to 0.15] for Sinovac versus 0.00 [IQR 0.00 to 0.01], both *P*<0.0001)(Fig. 2D, bottom). BioNTech elicited significantly higher proportions of spike-specific IFN-γ^+^CD4^+^T (*P*<0.0001) but not IFN-γ^+^CD8^+^T cells (*P*=0.6376) including all polyfunctional subsets (all *P*<0.0001) as compared with the Sinovac group (Fig. 2D and 2E). Spike-specific CD8^+^T cells were consistently detected in 81.8% of our BioNTech-vaccinees similar to recent reports by others (19–21). BioNTech, however, did not elicited significantly higher frequencies of spike-specific IFN-γ^+^CD8^+^T cells (*P*=0.6376) as compared with the Sinovac group (Fig. 2D and 2E). Surprisingly, Sinovac did not induce measurable levels of NP-specific IFN-γ^+^CD4^+^T or IFN-γ^+^CD8^+^T cells compared with the non-vaccinated group (Fig. 2D). As expected, CMV-specific IFN-γ^+^CD4^+^ and CD8^+^ T cells were comparable between these two vaccinated groups (Fig. S2A and S2B). The slightly lower levels of CMV-specific IFN-γ^+^CD4^+^ and CD8^+^ T cells in the BioNTech group than the non-vaccinated control (*P*=0.0457 and 0.0464, respectively) might indicate that there was unlikely vaccine-elicited bystander spike-specific T cell responses (Fig. S2A and S2B). In addition, spike-specific IFN-γ^+^CD4^+^ T cells induced by both vaccines showed similar phenotypic profiles of dominated effector memory subsets (Fig. 2F). These results demonstrated that while spike-specific CD4^+^T and CD8^+^T cells were generated by both vaccinees, Sinovac induced significantly weaker spike-specific CD4^+^T cell responses including polyfunctional subsets as compared with BioNTech.

### NAbs against SARS-CoV-2 VOCs among BioNTech and Sinovac vaccinees

Considering the rising issues of SARS-CoV-2 variants of concerns (VOCs) on the ongoing pandemic (7–10, 22), we tested plasma neutralization against multiple VOCs including B.1.1.7 (alpha), B.1.351 (beta), P1 (gamma), B.1.617.2 (delta) and B.1.429 (Epsilon). In general, the amounts of cross-NAbs against VOCs elicited by BioNTech were significantly stronger than those by Sinovac with 5-16-fold differences of the NAbTs including 6.72-fold against D614G (451.8 [95%CI 341.1 to 598.6] versus 67.15 [95%CI 38.15 to 118.2]), 7.09-fold against B.1.1.7 (593.5 [95%CI 422.7 to 833.2] versus 83.72 [95%CI 47.68 to 147.0]), 7.13-fold against B.1.351 (107.0 [95%CI 62.1 to 184.4] versus 15.0 [95%CI 9.24 to 24.37]), 15.73-fold against P1 (548.9 [95%CI 403.5 to 746.6] versus 34.9 [95%CI 20.01 to 60.9]), 5.64-fold against B.1.617.2 (131.0 [95%CI 91.56 to 187.3] versus 23.19 [95%CI 14.49 to 37.11]), and 10.36-fold against B.1.429 (565.8 [95%CI 425.7 to 752.0] versus 54.59 [95%CI 31.67 to 94.11]) (Fig. 3A) (all *P* values <0.0001) (Figure 3A). Based on NAbTs, study subjects were stratified into four groups: no response (<20: lower than LOQ), low (20-256), medium (256-1024), and high (>1024). We found that significantly fewer Sinovac vaccinees than BioNTech recipients had measurable NAbTs against VOCs, including 82.1% [23/28] against D614G (*P*=0.015), 85.7% [24/28] against B.1.1.7 (*P*=0.039), 14.3% [4/28] against B.1.351 (*P*<0.0001), 53.6% [15/28] against P1 (*P*<0.0001), 50.0% [14/28] against P1 (*P*<0.0001), and 75.0% [21/28]against B.1.429 (*P*=0.002). Moreover, most Sinovac responders showed low NAbTs. In contrast, only 7 (20.6%) and 2 out of 34 (5.9%) BioNTech vaccinees lacked NAbTs against B.1.351 and B. 1.617.2, respectively (Fig. 3B). Notably, NAbTs elicited by BioNTech were significantly reduced against VOCs relative to the wildtype virus including −3.1-fold against D614G, −2.4-fold against B.1.1.7, −13.09-fold against B.1.351, −2.55-fold against P1, −10.91-fold against B.1.617.2 and −2.52-fold against B.1.429 (all *P* <0.0001) (Fig. 3C). Similar NAbTs reduction against VOCs was also observed among Sinovac responders including −1.12-fold against D614G, −4.91-fold against B.1.351 (*P*<0.0001), −2.16-fold against P1 (*P*=0.0111), −3.18-fold against B.1.617.2 (*P*=0.0001) and - 1.38-fold against B1.429 (Fig. 3D). Unexpectedly, however, the highest amount of cross-NAbs against VOCs, especially B.1.351 and B.1.617.2, was observed in a single Sinovac-vaccinee (Fig. 3D). These results demonstrated that NAb positive rates and titers against VOCs, especially against the beta B1.351 and the delta B.1.617.2 strains, were lower among Sinovac-vaccinees than BioNTech-vaccinees.

**Figure 3.**
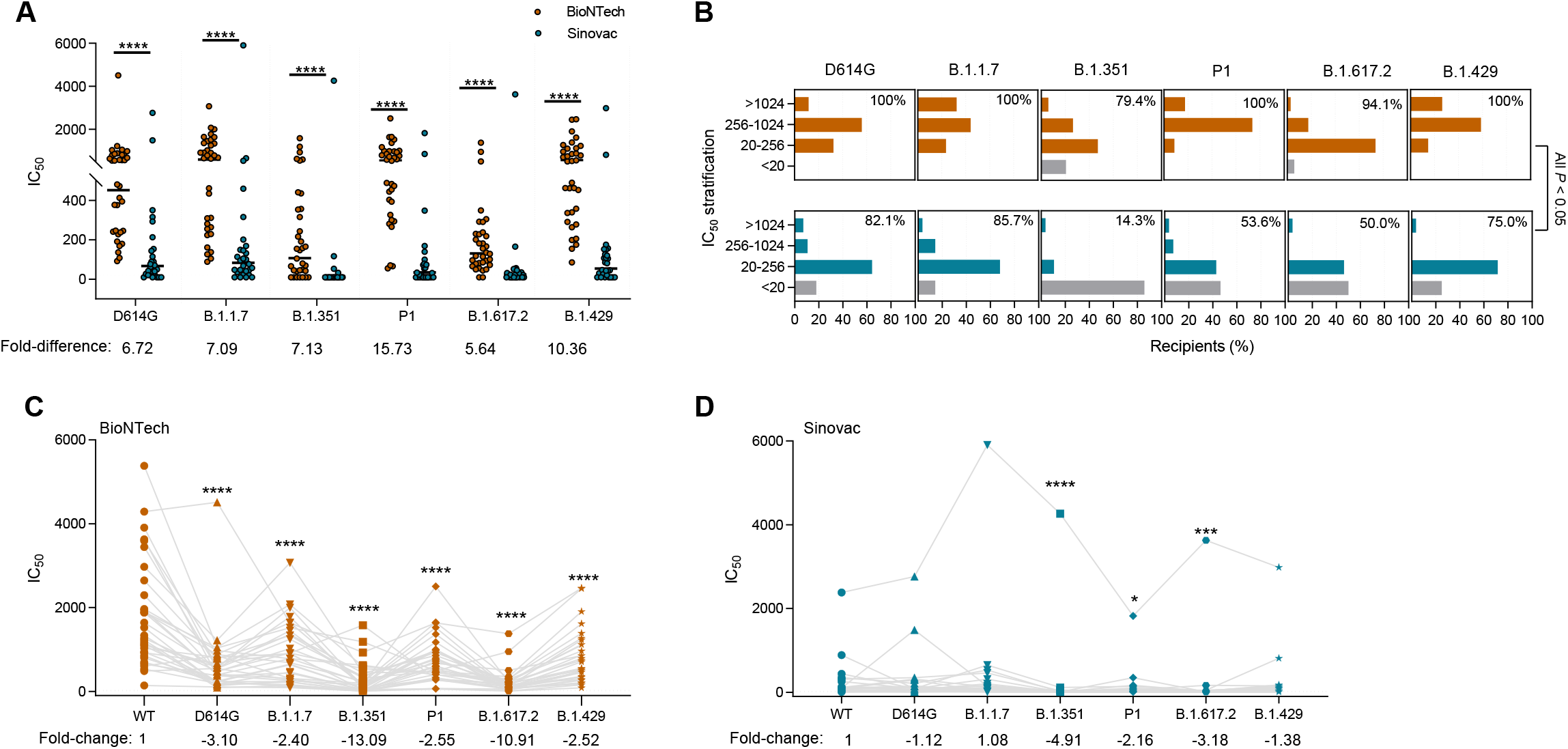
Neutralizing antibody activity against SARS-CoV-2 variants of concerns elicited by BioNTech and Sinovac. (**A**) Neutralizing antibody titers (NAbTs) against six SARS-CoV-2 strains from BioNTech (orange) (n=34) and Sinovac(green) (n=28) participants at acute phase after the second vaccination. NAbTs represent serum dilution required to achieve 50% virus neutralization (IC_50_). Numbers under the x-axis indicate the fold difference of BioNTech to Sinovac. (**B**) shown neutralizing IC_50_ of four response levels from BioNTech (orange) and Sinovac (green) recipients. Grey bars indicate the percentage of non-responders. Numbers in the top right corner represent the percentage of responders. Neutralizing IC_50_ against wild type compared to B.1.1.7, B.1.351, P1, B.1.617.2 and B.1.429 in BioNTech (**C**) and Sinovac (**D**) vaccinees. Numbers under the x-axis indicate the fold change of different VOC relative to wild type. Mann-Whitney U tests was used for between-group comparison in A, C and D. Two-sided chi-square tests were used in B. The bars represent geometric mean in A, C, D. Dotted black lines indicate the limit of quantification (LOQ. \**P*<0.05, \*\**P*<0.01, \*\*\**P*<0.001, \*\*\*\**P*<0.0001.

### Vaccine-induced NAbs against VOCs and T cell responses in the memory phase

Since the epidemic was well controlled in Hong Kong, no SARS-CoV-2 infection was found among our study subjects at the follow-up time of 3 months after the second vaccination. We were able to follow-up 43 longitudinal subjects to measure their immune responses at the memory phase. There is a −1.92-fold reduction of anti-spike IgG (*P*=0.0131) in the Sinovac group (Figure 4A) while a −1.31 fold reduction of anti-RBD IgG (*P*=0.0229) was found in the BioNTech group (Figure 4B). Surprisingly, there were significant reductions of NAbTs against the wildtype in both BioNTech-vaccinees (−2.72 fold, *P*<0.0001) and Sinovac-vaccinees (−4.03 fold, *P*=0.0001)(Figure 4C). Similar reductions were found against the panel of VOCs among BioNTech- and Sinovac-vaccinees including D614G (−4.68 fold, *P*<0.0001 and −5.32 fold, *P*=0.0084) (Figure 4D), B.1.1.7 (−4.38 fold, *P*<0.0001 and −5.14 fold, *P*=0.0002) (Figure 4E), B.1.351 (−2.22 fold, *P*=0.0003 and no difference) (Figure 4F), P1 (−6.33 fold, *P*<0.0001 and −3.06, *P*=0.0017) (Figure 4G), B.1.617.2 (−3.38 fold, *P*<0.0001 and - 1.53 fold, *P*=0.0078) (Figure 4H) and B.1.429 (−2.52 fold, *P*<0.0001 and no difference) (Figure 4I), respectively. In terms of spike-specific T cell responses, a significant decrease of spike-specific IFN-γ^+^CD4^+^T cells at the memory phase was observed for both vaccine groups (−2.69 fold, *P*<0.0001 and −2.4 fold, *P*=0.029), respectively (Figure 4J). A similar decrease of spike-specific IFN-γ^+^CD8^+^T cells at the memory phase was mainly observed for BioNTech-vaccinees (−2.94 fold, *P*<0.0001). A similar trend was found with Sinovac-vaccinees but without statistical significance and one individual showed an big increase of IFN-γ^+^CD8^+^T cells in the memory phase (Figure 4K). These results demonstrated that both vaccine-induced NAb and T cell responses have waned significantly at the memory phase just three months after the second immunization. In particular, since most Sinovac-vaccinees have low or unmeasurable NAbTs at this stage, they may face higher risk to infection by the spreading VOCs.

**Figure 4.**
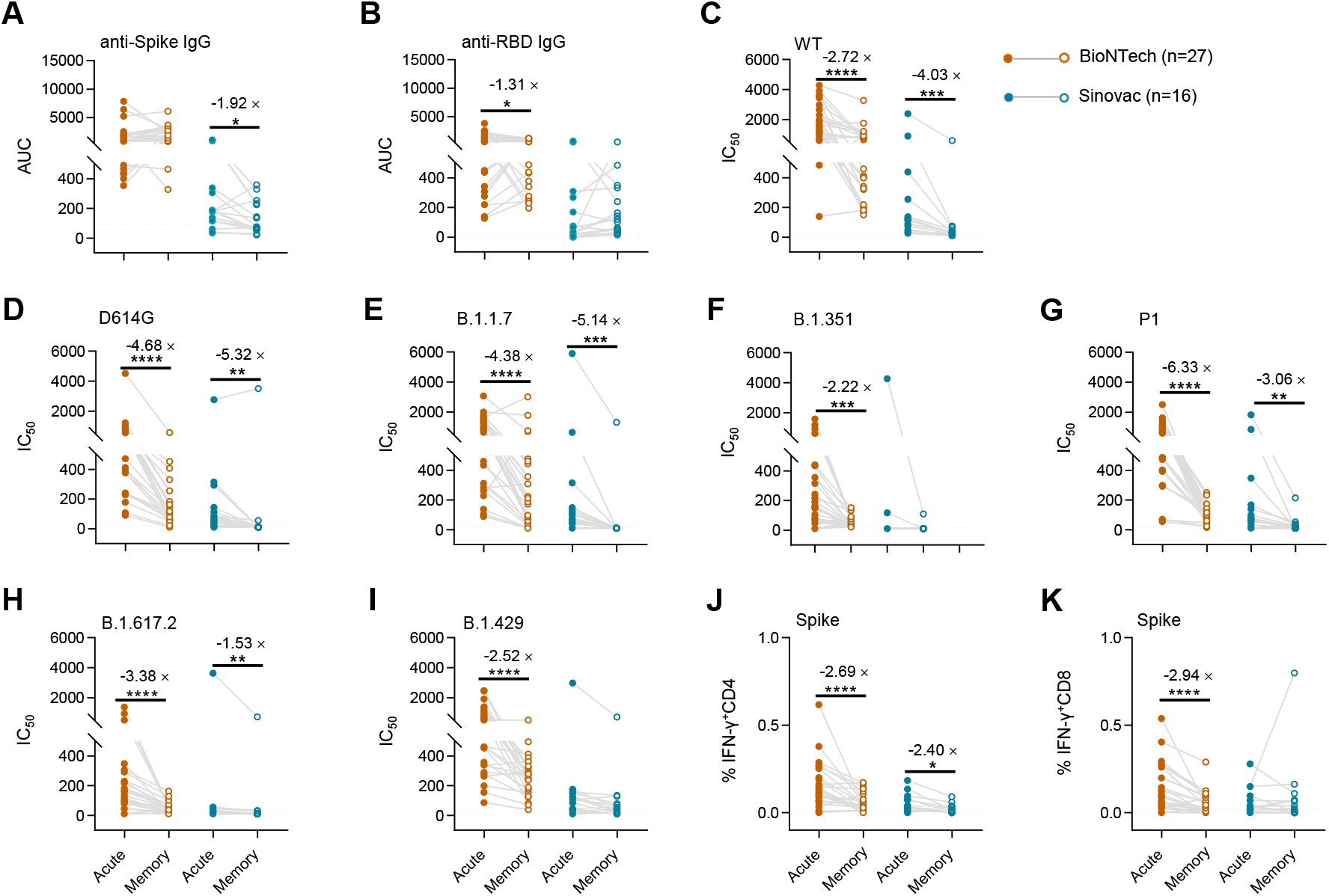
Changes of humoral and cellular responses from the acute phase to the memory phase among BioNTech- and Sinovac-vaccinees. Longitudinal samples from 27 BioNTech and 16 Sinovac vaccinees were available to track immune response from acute phase to memory phase. Longitudinal binding antibodies to anti-Spike (**A**) and RBD IgG (**B**), neutralizing IC_50_ to wild type (**C**) and different VOCs including D614G (**D**), B.1.1.7 (**E**), B.1.351 (**F**), P1 (**G**), B.1.617.2 (**H**), B.1.429 (**I**) and spike-specific IFN-γ^+^ CD4^+^ (**J**) or CD8^+^ (**K**) T cells were measured and compared. Significant differences between acute phase and memory phase of both vaccine group were determined by Wilcoxon signed-rank. Dotted black lines indicate the limit of quantification (LOQ. \**P*<0.05, \*\**P*<0.01, \*\*\**P*<0.001, \*\*\*\**P*<0.0001.

### Correlation analysis of vaccine-induced humoral and cellular immune responses

Considering that NAbTs correlate with viral infectivity (23), we conducted similar correlation analysis at acute phase. Similar to the findings described previously by us (16) and others (19, 24), strong positive correlations were found between NAbTs against wildtype pseudovirus and Spike-specific IgG (Figure 5A, r=0.7756 and *P*<0.0001) or RBD-specific IgG (Figure 5B, r=0.8241 and *P*<0.0001). Furthermore, strong positive correlations were found between NAbTs against wildtype and NAbTs against D614G (Figure 5C, r=0.8314 and *P*<0.0001) or NAbTs against B.1.1.7 (Figure 5D, r=0.8426 and *P*<0.0001) or NAbTs against B.1.351 (Figure 5E, r=0.7381 and *P*<0.0001) or NAbTs against P1 (Figure 5E, r=0.8902 and *P*<0.0001), or NAbTs against B.1.617.2 (Figure 5G, r=0.7591 and *P*<0.0001) or NAbTs against B. 1.429 (Figure 5H, r=0.8761 and *P*<0.0001). These results indicated that NAbTs against wildtype virus at peak immunity predicted NAbs cross-reactivity despite the titer drops against these VOCs tested. In addition, NAbTs against wildtype was correlated with the frequency of spike-specific IFN-γ^+^CD4^+^T cell response (Figure 5I, r=0.5805 and *P*<0.0001) but not with the frequency of spike-specific IFN-y^+^CD8^+^T cell frequencies (Figure 5J, *P*=0.9482). These results indicated that CD4 helper T cells had likely contributed to the induction of NAbs.

**Figure 5.**
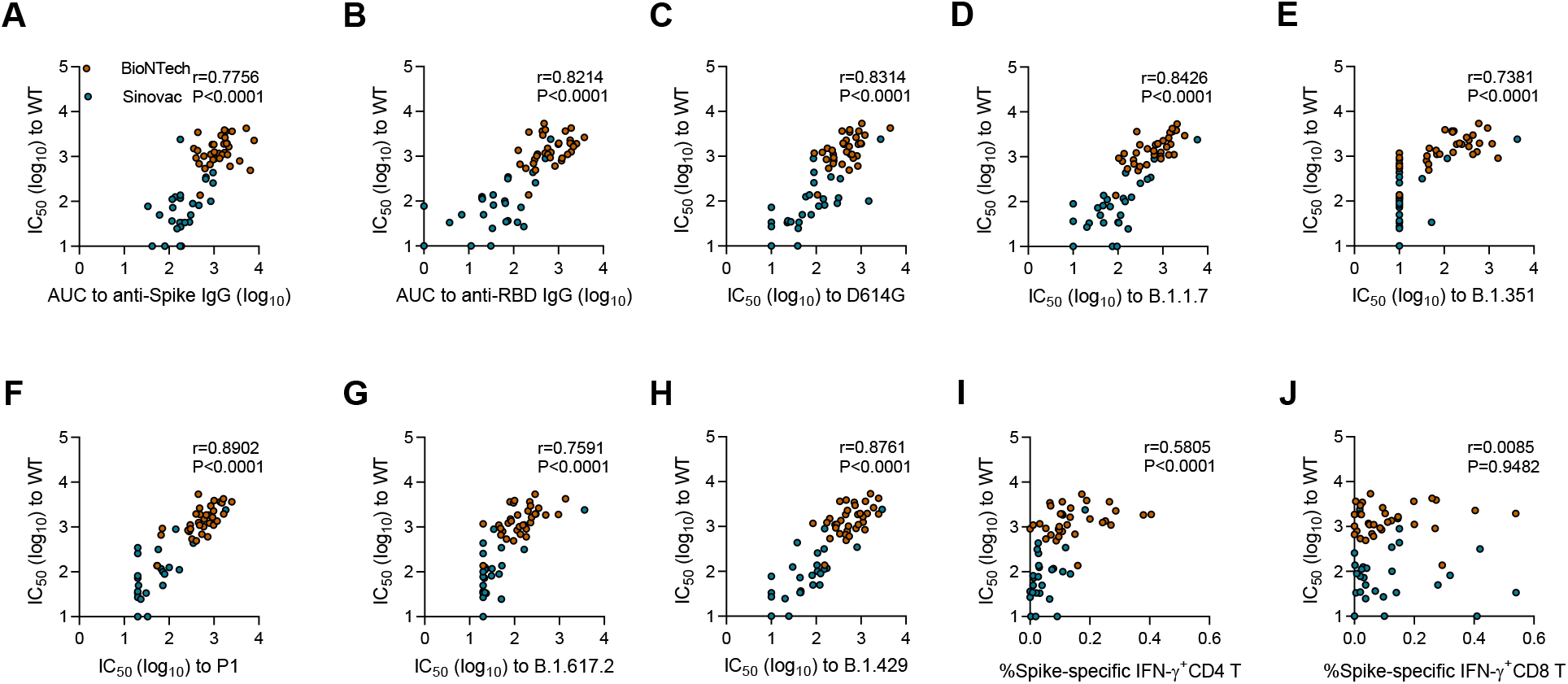
Correlation analysis of vaccine-induced humoral and cellular immune responses. Correlation analysis of anti-Spike (**A**) and RBD IgG (Iog10) (**B**), IC_50_ against different VOCs including D614G (**C**), B.1.1.7 (**D**), B.1.351 (**E**), P1 (**F**), B.1.617.2 (**G**),B.1.429 (**H**) and spike-specific IFN-γ^+^ CD4^+^ (**I**) or CD8^+^ T cells (**J**) to IC_50_ against wild type (WT). The non-parametric Spearman test was used for correlation analysis. \**P*<0.05, \*\**P*<0.01, \*\*\**P*<0.001, \*\*\*\**P*<0.0001.

## Discussion

Here we report a prospective longitudinal study of antibody and T cell immune responses among BioNTech- and Sinovac-vaccinees in Hong Kong. To the best of our knowledge, our results present the first clinical study on NAb responses against the global panel of VOCs and T cell responses to wildtype induced by the standard 2-dose BioNTech and Sinovac vaccinations in parallel. Both vaccines were safe and well-tolerated among our study subjects although BioNTech induced more frequent but transient side effects. While both vaccines induced NAb and S-specific T cell responses to the wildtype virus similar to previous findings (2, 6, 25), the geometric NAbTs of Sinovac-vaccinees was 19-fold lower than that of BioNTech-vaccinees (73.7 versus 1400.5), which is consistent to recent publications (26, 27). The NAb response corelated positively with CD4 but not CD8 responses, suggesting that vaccine-induced CD4 helper probably contributes to B cell activation. Moreover, our findings on waning NAb responses to wildtype virus among BioNTech-vaccinees are consistent to several studies published recently (19, 24, 28, 29). Importantly, against the global panel of VOCs, NAb response rates and titers among Sinovac-vaccinees were not only significantly lower but also disappeared more dramatically just three months after the vaccinations as compared with BioNTech-vaccinees. Sinovac-vaccinees also exhibited lower neutralization potency index and waning S-specific IFN-γ^+^CD4^+^T cells while they showed no advantage for inducing S-specific IFN-γ^+^CD8^+^T cells. The neutralization potency index is predictive of immune protection against symptomatic SARS-CoV-2 infection (30). Our findings are in line with better clinical efficacy of BioNTech than that of Sinovac during phase 3 trials against COVID-19 (95.0% versus 50.65% - 83.5% based on Sinovac trials from Brazil, Turkey and Indonesia)(1, 5, 31). Since Sinovac is one of the most extensively used vaccine with approaching 1.9 billion doses administrated in many countries, our findings have significant implication to Sinovac-vaccinees who may face higher risk than BioNTech-vaccinees to the spreading VOCs breakthrough infection and should be considered as a priority for the third vaccination.

SARS-CoV-2 VOCs continue to emerge globally, which have brought new challenges to the efficacy of COVID-19 vaccines in emergency use (7–10) (22). The B.1.351 variant escaped not only RBD-specific monoclonal NAbs but also vaccine-induced NAbs and convalescent sera (7). Similarly, we found that our subjects contained the lowest NAbTs to B.1.351 in sera elicited by both BioNTech and Sinovac. In particular, the NAbTs induced by BioNTech to B.1.351 decreased by 13-fold, which is worse than the 6.5-8.6-fold decrease among American vaccinees (7). In this study, while 79.4% BioNTech-vaccinees developed NAbs to B.1.351, only 14.% Sinovac-vaccinees had similar NAbs. Fortunately, B.1.351 and its variants have not becoming a major circulating VOC globally. Instead, the B.1.617.2 variant has become the major VOC after its first detection at the end of March 2021 in Indian (32). Due to its extremely high transmissibility and infectivity, cases of breakthrough B.1.617.2 infections have been increasing dramatically even in regions with high vaccination coverage (33). We found that 94.1% BioNTech-vaccinees and 50% Sinovac-vaccinees have developed cross-NAbs to B.1.617.2 mainly at low NAbTs (20-256). Compared with NAbTs to the wildtype, there were 10.91-fold and 3.18-fold reduced NAbTs observed for BioNTech- and Sinovac-vaccinees, respectively, in line with the 5.8-fold decrease against B.1.617.2 induced by mRNA vaccine in the UK and other studies (22, 34). Since the geometric mean NAbTs further dropped by 3.38- and 1.53-fold just three months after the vaccination, especially with most Sinovac-vaccinees to the detection limit (<20), the efficacy of preventing breakthrough B.1.617.2 infections is indeed worrisome. Nevertheless, the efficacy of 2-dose vaccinations against B.1.617.2 was 88% for BioNTech and 67% for ChAdOx1 nCoV-19 (35), while the efficacy of 2-dose inactivated vaccinations was 59% in a test-negative case-controlled study (36). Recently, an exploratory trial of boosting with the third dose of inactivated SARS-CoV-2 showed induction of 7.2-fold higher NAbTs, together with 5.9- and 2.7-fold higher spike-specific CD4^+^ and CD8^+^ T cells one week after the third dose (37). The third timely boost vaccination is likely helpful especially for Sinovac-vaccinees against breakthrough infections against the pandemic VOCs. Having said so, we urge that populations that have not even received the complete 2-dose vaccinations should be given the highest priority to reduce cases, hospitalizations and deaths.

This study has some limitations. Due to lack of a spreading epidemic in Hong Kong, we could not determine vaccine-mediated protective efficacy. The sample size of this study was relatively small and most of our subjects have not reached 6 months for follow-up testing. Extend the follow-up to 6 months and one year or longer is necessary for future study. While NAbs have been indicated as correlates of protection (23, 30), the protective role of vaccine-induced T cell responses remains to be further investigated. During acute SARS-CoV-2 infection, we and others demonstrated that antigen-specific T cell responses have likely been associated with viral control and limited pathogenesis(13) (38). In this study, while we consistently found antigen-specific CD4^+^T cells after vaccinations by both types of vaccines as previously reported by others (19, 39, 40), majority of our subjects did not show measurable RBD-specific CD4^+^ T cells. The difference between spike- and RBD-specific CD4 responses and why only spike-specific CD4 responses correlated to NAbTs but not to CD8^+^ T cells remain unclear. VOC spike-specific T cell responses were not explored due to limited cells received, although some studies indicated that the mutations in VOCs might modify single T cell specificities but could not fully escape the whole repertoire of spike-specific T cells (41, 42). Future studies are needed to address these limitations.

## Supporting information

Supplemental meterials

## Contributors

Z.C. supervised the collaborative team, conceived of and designed the study, and wrote the manuscript. K.M. coordinated donor recruitment and specimen collection. Q.P., R.Z., Y.W. designed some experiments, analysed the data, and prepared the manuscript. M.Z., N.L., S.L., H. H., K.A. and D.Y. performed immune assays, H.W. and K.Y.Y. provided critical comments, supports and materials.

## Acknowledgements

We thank Dr. David D. Ho for kindly providing the expression plasmids encoding for D614G, B.1.1.7 and B.1.351 variants and Dr. Linqi Zhang for B. 1.617.2.

## Data Sharing Statement

The authors declare that the data supporting the findings of this study are available from the corresponding author upon request.

---

**Figure S1. Flow cytometry analysis of antigen-specific T cell response.** (**A**) Gating strategies to define CD3^+^CD4^+^ and CD3^+^CD8^+^T cells. (**B**) Representative dot plots depict IFN-γ^+^, TNF-α^+^ and IL-2^+^CD4^+^ and CD8^+^ T cells after stimulated with SARS-CoV-2 RBD, Spike, NP as well as CMV and negative control (CD28CD49d only). Representative examples are from a BioNTech and Sinovac vaccinee, respectively.

**Figure S2. CMV-specific T cell responses in vaccinees and non-vaccinated subjects.** Comparison of CMV-specific IFN-γ^+^CD4^+^ (**A**) and CD8^+^ T cells (**B**) among BioNTech (n=33), Sinovac vaccinees (n=28) and non-vaccinated subjects (n=15). Responder rate were depicted under x-axis. Mann-Whitney U tests was used for between-group comparison. \**P*<0.05.

