## Supplemental meterials for "Waning immune responses against SARS-CoV-2 among vaccinees in Hong Kong": Supplemental Figures .pdf

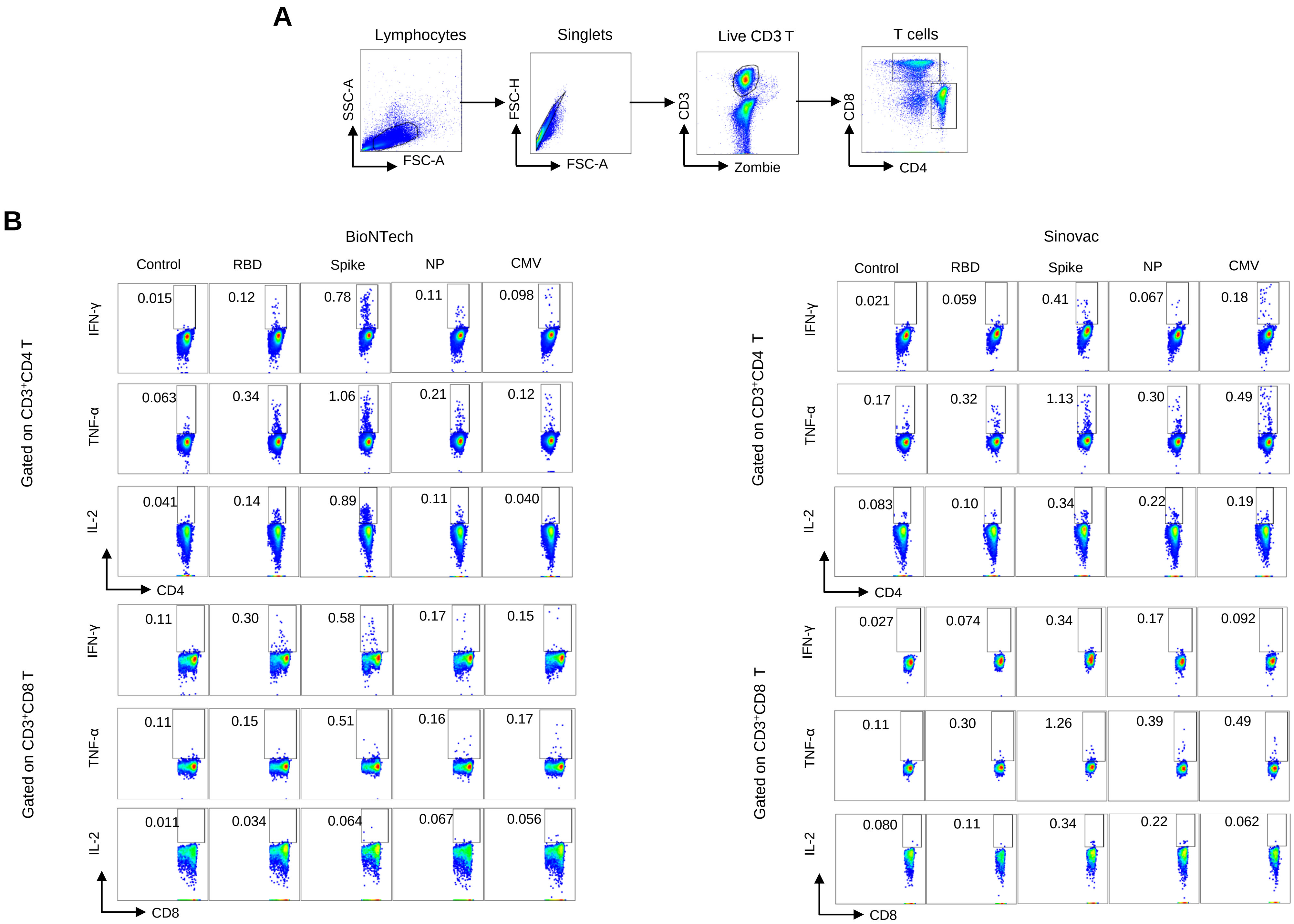
A
Lymphocytes
Singlets
Live CD3 T
T cells
SSC-A
FSC-A
FSC-H
FSC-A
CD3
Zombie
CD8
CD4
B
BioNTech
Sinovac
Control
RBD
Spike
NP
CMV
Gated on CD3+CD4 T
IFN-γ
TNF-α
IL-2
CD4
0.015
0.12
0.78
0.11
0.098
0.063
0.34
1.06
0.21
0.12
0.041
0.14
0.89
0.11
0.040
0.021
0.059
0.41
0.067
0.18
0.17
0.32
1.13
0.30
0.49
0.083
0.10
0.34
0.22
0.19
Gated on CD3+CD8 T
CD8
0.11
0.30
0.58
0.17
0.15
0.11
0.15
0.51
0.16
0.17
0.011
0.034
0.064
0.067
0.056
0.027
0.074
0.34
0.17
0.092
0.11
0.30
1.26
0.39
0.49
0.080
0.11
0.34
0.22
0.062

Figure S1

**Figure S1. Flow cytometry analysis of antigen-specific T cell response.** (**A**) Gating strategies to define $CD3^+CD4^+$ and $CD3^+CD8^+$T cells. (**B**) Representative dot plots depict IFN-$\gamma^+$, TNF-$\alpha^+$ and IL-$2^+CD4^+$ and $CD8^+$ T cells after stimulated with SARS-CoV-2 RBD, Spike, NP as well as CMV and negative control (CD28CD49d only). Representative examples are from a  BioNTech and Sinovac vaccinee, respectively.

### Figure S2

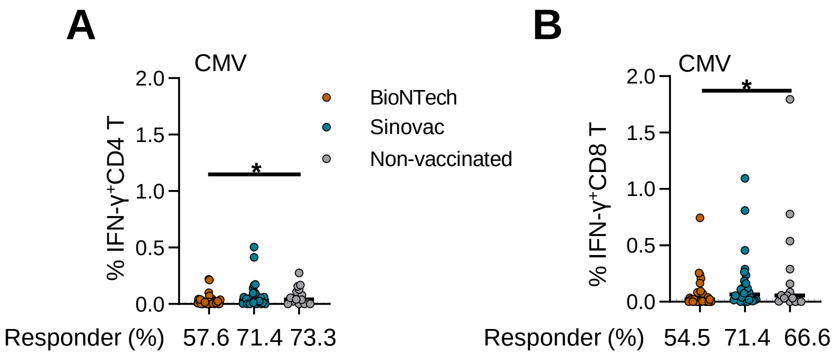


**Figure S2. CMV-specific T cell responses in vaccinees and non-vaccinated subjects.** Comparison of CMV-specific IFN-$\gamma^{+}$CD4$^{+}$ (**A**) and CD8$^{+}$ T cells (**B**) among BioNTech (n=33), Sinovac vaccinees (n=28) and non-vaccinated subjects (n=15). Responder rate were depicted under x-axis. Mann-Whitney U tests was used for between-group comparison. *$P$<0.05.
